# Versatile and Portable Cas12a-mediated Detection of Antibiotic Resistance Markers

**DOI:** 10.1101/2024.11.14.623642

**Authors:** Maryhory Vargas-Reyes, Roberto Alcántara, Soraya Alfonsi, Katherin Peñaranda, Dezemona Petrelli, Roberto Spurio, Monica J. Pajuelo, Pohl Milon

**Author notes:** Correspondence: Pohl Milon; Roberto Alcántara.

## Abstract

Antibiotic-resistant bacteria are spreading in clinical, industrial, and environmental ecosystems. The spreading dynamics to and from the environment are unknown, largely due to the lack of appropriate (robust, fast, low-cost) analytical assays. In this study, we developed C12a, a versatile molecular toolbox to detect genetic markers of antibiotic resistance using CRISPR/Cas12a. Biochemical characterization show that the C12a toolbox can detect less than 100 attoMolar of pure DNA fragments from the *blaCTX-M15* and *floR* genes, conferring resistance to b-lactams and amphenicols, respectively important for human and veterinary uses. In microbiological assays, C12a detected less than 10^2^ CFU/mL and high concordance was observed if compared to antibiotic susceptibility tests, PCR, or to whole genome sequencing. Additionally, C12a confirmed a high prevalence of the integrase/integron system in *E. coli* isolates containing multiple antibiotic resistance genes (ARGs). The C12a toolbox shows equivalent detection performance in diverse laboratory settings, results redout (Fluorescence *vs* FLA) or input sample. Altogether, this work presents a comprehensive proof-of-concept, development description, and biochemical characterization of a collection of molecular tools to detect antibiotic resistance markers in a one health setup.

## Introduction

The discovery of antibiotics allowed control of bacterial infections, improving surgical outcomes, treating infectious diseases, and boosting farm productivity, ultimately contributing to increased life expectancy [1]. However, the antimicrobial resistance (AMR) threat accelerated due to the combination of multiple interconnected factors, including antibiotic overuse[2], [3], complex ecological factors [4], [5], ineffective waste management and mechanisms of spread by horizontal gene transfer [6]. Effective surveillance strategies are lacking due to insufficient capacities for sensitive, timely, cost-effective, and accurate monitoring [7], [8]. Among the most concerning mechanisms are Extended Spectrum β-Lactamases (ESBLs) and resistance to amphenicols, due to their relevance for both human and animal health [9]. In Gram-negative bacteria, the *blaCTX-M-15* and *floR* are two of the most frequently reported antibiotic resistance genes (ARGs) within their respective classes, conferring resistance to β-lactam and amphenicol antibiotics, respectively (Figure 1A-B) [10]. While *blaCTX-M* [11], [12] variants have been primarily associated with bacterial human isolates, *floR*[13], [14] has traditionally been linked to animal-derived bacterial isolates, as florfenicol is broadly used in livestock production. However, bacteria *floR*-positive strains are increasingly detected in humans, particularly among individuals in close contact to poultry [15]. Both ARGs have been detected in bacteria isolates from clinical[16], [17], [18], [19], community-acquired[12], [15], animal[13], [14], [20], [21], and environmental samples[10] including water[22], [23] and farms soil[23]. The ARGs spread is enhanced by mobile genetic elements (MGEs), due to their ability to move within and between different host species [24]. Among them, integrons are genetic elements that mobilize and express gene cassettes that confer resistance to antibiotics, disinfectants, heavy metal, and other pollutants [25]. Although integrons are not mobile on their own, they are frequently integrated within transposons or carried on plasmids, facilitating their horizontal transfer [26]. Class 1 integrons are the most prevalent in both clinical[27] and environmental [26], [28], [29] contexts, particularly in *Enterobacteriaceae* clinical strains, such as *E*.*coli* [30] (Figure 1C). Its relative abundance has been proposed as a marker of anthropogenic AMR pressure [31], since genomic analyses of cassette arrays frequently show co-occurrence with resistance to aminoglycoside, sulfonamide, tetracycline, β-lactams, and amphenicol [32], [33]. Due to its widespread distribution and frequent co-occurrence with multiple ARGs, detection of Class 1 integron could be used as a proxy to assess the ecological impact of human activity in AMR surveillance [25], [32].

**Figure 1.**
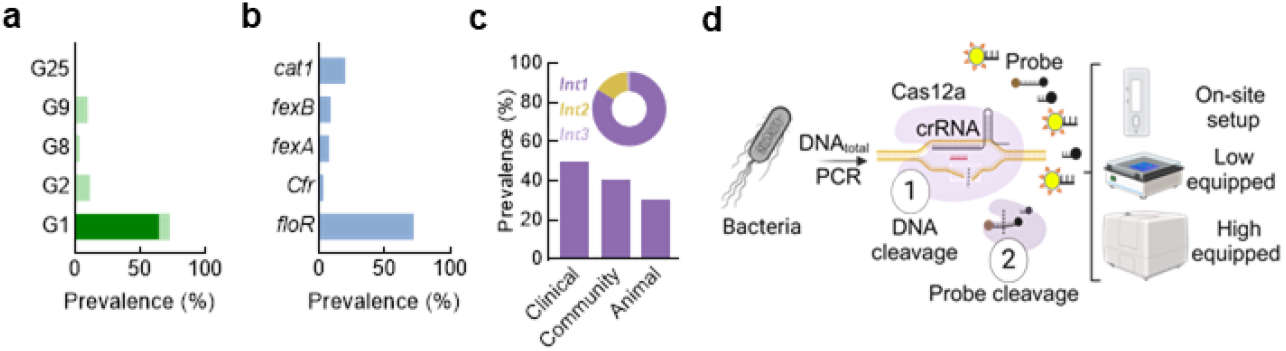
*blaCTX* and *floR* epidemiological context and the C12a rationale. **(A)** Prevalence and diversity of beta-lactamases in *E. coli* [11], [12]. The darker green bar represents the prevalence of the *blaCTX-M-15* gene within the Group 1. **(B)** Prevalence and diversity of amphenicol resistance genes in bacteria [13], [14]. **(C)** Prevalence of the Class 1 Integron across clinical [30], community, and animal settings [39]. The pie chart shows the three integron classes distribution in environmental isolates [40] **(D)** Scheme showing the C12a toolbox and the molecular mechanism of Cas12a-mediated detection, highlighting signal development, versatility, and portability.

Currently, different methods for detecting antimicrobial resistance (AMR) are used, with the most common being antimicrobial susceptibility testing (AST) and molecular techniques such as PCR, qPCR, and DNA sequencing [34], [35]. While these methods are considered standards, there is a need of assays with new technologies that combine the velocity provided by molecular methods with the low-costs of AST, and applicability in resource-limited or on-site settings. Target pre-amplification followed by nucleic acid detection using CRISPR technology has shown promising results due to the high specificity, sensitivity, low costs, and speed of the assay [36]. The molecular detection mechanism involves crRNA, a targeting sequence homologous to the gene of interest. crRNA hybridization with the complementary target region activates the Cas12a endonuclease, which then engages in trans-cleaving activity, cutting ssDNA non-specifically [37]. When using a fluorescence donor/quencher FRET pair placed into a ssDNA, this trans-cleaving activity generates an increase in fluorescence signal that can be detected using a wide range of laboratory set ups [38](Figure 1D).

Here, we present the C12a toolbox, a CRISPR-based set of molecular tools designed to detect genetic elements that confer antibiotic resistance or contribute to ARG propagation. The toolbox includes assays detecting *blaCTX-M-15* (C12a^bCTX^), *floR* (C12a^FLO^), and the *intI1* integrase gene (C12a^INT^), selected for their relevance within the One Health framework. We describe the proof-of-concept, the biochemical and analytical characterization of C12a as an adaptable platform optimized for high-, and low-equipped laboratories in addition to an on-site portable setting (Figure 1C). Implementation of C12a nucleic acid-based technologies could effectively support fast, accurate, and sensitive detection, contributing to ARG surveillance efforts.

## Results

### C12a Development

ARG detection sites were defined by comparing 61 *blaCTX-M-15* and 46 *floR* sequences from *E. coli* isolates available in GenBank [41] (Supplementary Dataset 1 and Dataset 2, respectively). Based on consensus sequences two primer sets for *blaCTX-M-15* and one for *floR* were selected to amplify regions containing a PAM sequence for Cas12a (TTTN) (Figure 2A, Figure S1, Table S2). Agarose gel electrophoresis confirmed amplification of *blaCTX-M-15* (216 bp) and *floR* (270 bp), with no bands in negative controls (Figure S2). Amplicons were sequenced to confirm the presence of the expected target DNA for each gene (Table S3). To optimize the crRNA/Cas12a reaction, two candidate crRNAs for each target gene were designed to contain a complementary sequence to the amplified fragments (Table S2). To determine sensitivity parameters, an initial test with increasing Mg^2+^ concentrations revealed an optimal value of 20 mM, which was used in all assays. Both *blaCTX-M-15* and *floR* amplicons were recognized by crRNA/Cas12a complex resulting in an increase in fluorescence signal (Figure S3) For *blaCTX-M-15* target gene, primer set 2 combined with crRNA^CTX-2^, showed greater fluorescence increase compared to crRNA^CTX-1^. For *floR* gene, crRNA^FLO-1^ outperformed crRNA^FLO-2^ (Figure S1). Thus, crRNA^CTX-2^ and crRNA^FLO-1^ were selected for further assays. Regardless of template concentration, crRNA^CTX^ consistently generated a fluorescence signal faster than the *floR* crRNAs (Figure 2B and Figure S1).

**Figure 2.**
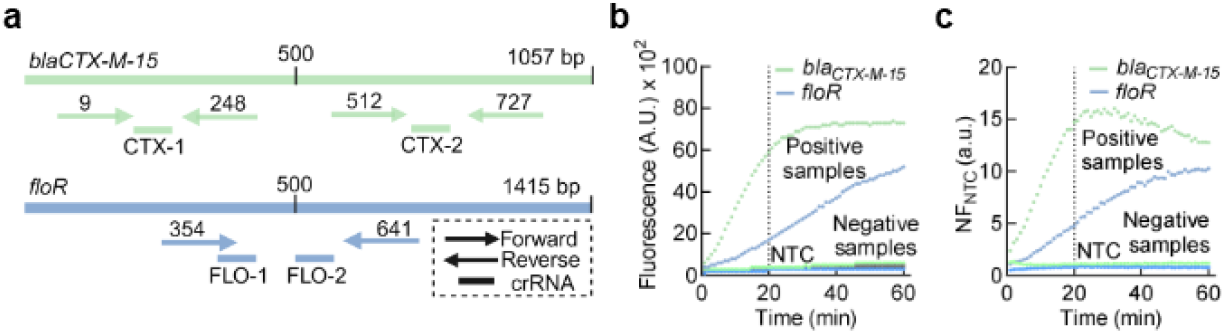
Experimental setup for CRISPR/Cas12a-based assay development. **(A)** Primers and crRNA location on *blaCTX-M-15* and *floR* gene targets, ensuring coverage of PAM regions and complementary crRNA areas. **(B)** Example of raw fluorescence readouts from CRISPR/Cas12a assays, showing a time-dependent increase of the signal. Positive and negative controls, alongside the Non-Template Control (NTC), are represented to illustrate the assay’s ability to differentiate samples with or without ARGs. **(C)** Example of normalized fluorescence ratio NF_NTC_ (a.u.) over time. In **(B)** and **(C)**, the dotted line indicates the 20-minute detection point, selected as the cut-off reading time for analysis.

The results of this assay are reported as fluorescence ratio to the no-template control (NTC), indicated as NF_NTC_ (a.u.). To calculate NF_NTC_, each raw fluorescence value was divided by the corresponding value from the NTC (Figure 2C). For the *blaCTX-M-15* gene, positive samples exhibited an NF_NTC_ of at least 14-fold, while negative samples display NF_NTC_ values near 1. Similarly, for the *floR* gene, positive samples showed a 4-fold increase in NF_NTC_ compared to negative controls (Figure 2C and Figure S1).

Based on performance, the final detection systems were defined. For *blaCTX-M-15* detection, the C12a^bCTX^ system incorporates the primer set 2 and crRNA^CTX-2^ (Table S2). On the other hand, Cas12a^FLO^ detects *floR* using the primer set 1 and the crRNA^FLO-1^ (Table S2).

### C12a Analytical Sensitivity

Two key analytical parameters describe any given assay for analyte detection: the limit of blank (LoB; also referred to as the cut-off) and the limit of detection (LoD). The average NF_NTC_ values for negative samples using C12a^bCTX^ or C12a^FLO^ were 1.04 ± 0.02 and 0.94 ± 0.02 after 20 minutes of incubation (Figure S4). Here we set the LoB as the average of true negative assays plus two standard deviations. Thus, the resulting LoB values were 1.08 NF_NTC_ (a.u.) for C12a^bCTX^ and 0.98 NF_NTC_ (a.u.) for C12a^FLO^. Based on these values, we can state that 95.45% of the values below the determined LoB were likely to be classified as nonspecific background signals (i.e., signals in the absence of the analyte) (Figure S4).

To estimate the LoD for both detection systems, purified and quantified amplicons were tested at increasing concentrations (from 10^−18^ to 10^−8^ M, atto to nanomolar). Averaged NF_NTC_ (a.u.) values, measured at 10-minute intervals were used to define the lowest detectable target concentration at the shortest reaction time. The LoD was calculated for each time interval, allowing the estimation of LoD as a function of reaction time (Figure 3A). The time dependence indicates that after 21 minutes for C12a^bCTX^ and 17 minutes for C12a^FLO^, the LoD reached half of its maximum value, suggesting that a 20-minute reaction time is a good compromise between assay speed and analytical sensitivity. The calculated LoD for C12a^bCTX^ and C12a^FLO^ at 20 minutes was 70 aM and 50 aM, respectively (Figure 3B). Taking NF_NTC_ values at times shorter than 20 minutes (i.e. 10 min) would result in a tenfold loss in sensitivity (Figure 3A).

**Figure 3.**
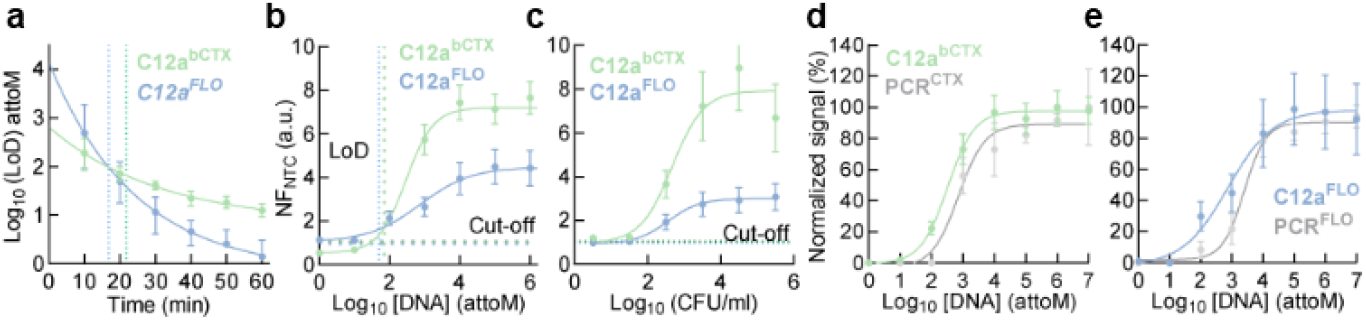
C12a^bCTX^ and C12a^FLO^ Limit of Detection (LoD). **(A)** Time-resolved determination of LoD values for both ARGs Cas12a reactions, plotted on a logarithmic scale. The graph illustrates the LoD decrease for C12a^bCTX^ and C12a^FLO^ over incubation time, with dotted lines marking the time-point when the LoD reaches half of its maximum value. **(B)** Normalized fluorescence ratio as a function of target concentration for ARG detection assays. The vertical dotted lines represent the LoD value calculated at 20 minutes, while the horizontal dotted lines represent NF_NTC_ LoB (cutoff). **(C)** Normalized fluorescence ratio over bacterial cell concentration for both C12a assays. The dotted horizontal lines indicate average NF_NTC_ cut-off values of 1.08 and 0.98 for C12a^bCTX^ and C12a^FLO^, respectively. **(D)** Normalized assay signals across target concentrations for C12a^bCTX^ detection compared to PCR-based detection evaluated by gel electrophoresis. The fluorescent ratio for the Cas12a reaction and electrophoresis gel band intensity, recorded as Area Under the Curve (AUC) values, are normalized as percentages to the highest value. **(E)** Same as (D), but for C12a^FLO^. Error bars in all panels represent standard deviations from 3 replicates.

We then assessed the analytical sensitivity of the CRISPR/Cas12a system in terms of the number of cells carrying the resistance genes (Figure 3C). Bacterial cultures of true positive *E. coli* ranging from 3 to 3×10^6^ CFU/mL were used (Figure S5). Non-linear fitting of the NF_NTC_ dependence on cell number (CFU) allowed the calculation of the LoD. C12a^bCTX^ reliably detected the *blaCTX-M-15* gene at 77 CFU/mL, while C12a^FLO^ required 173 CFU/mL to detect *floR* (Figure 3C). The lowest target concentration that could be empirically detected for the two target genes, was 10 to 100 times lower compared to gel electrophoresis analysis of PCR amplification (Figure 3D, E, and Figure S6). Altogether, the detection and sensitivity performance of C12a^bCTX^ and C12a^FLO^ provide a solid foundation for testing the detection systems on a larger number of samples (see below).

### ARG detection in *E. coli* isolates by C12a^bCTX^ and C12a^FLO^

The presence of antibiotic resistance genes (ARGs) and the related phenotypic resistance in *E. coli* isolates was assessed by two distinct assays: (i) molecular detection using the C12a toolbox and (ii) antimicrobial susceptibility testing (AST) (Figure 4). Two small independent sample sets were randomly selected from a collection of *E. coli* isolated from human sources in community settings (see Materials and Methods). Fifteen isolates were evaluated for the presence of *blaCTX-M-15* and beta-lactam resistance, while seventeen were assessed for *floR* and resistance to amphenicols. Phenotypic profiling was confirmed using the combined disk (CD) method for ESBL detection and broth microdilution to determine minimum inhibitory concentrations (MICs) for florfenicol (Table S4-5). The C12a^bCTX^ and C12a^FLO^ assays were then applied to their corresponding set to evaluate detection performance.

**Figure 4.**
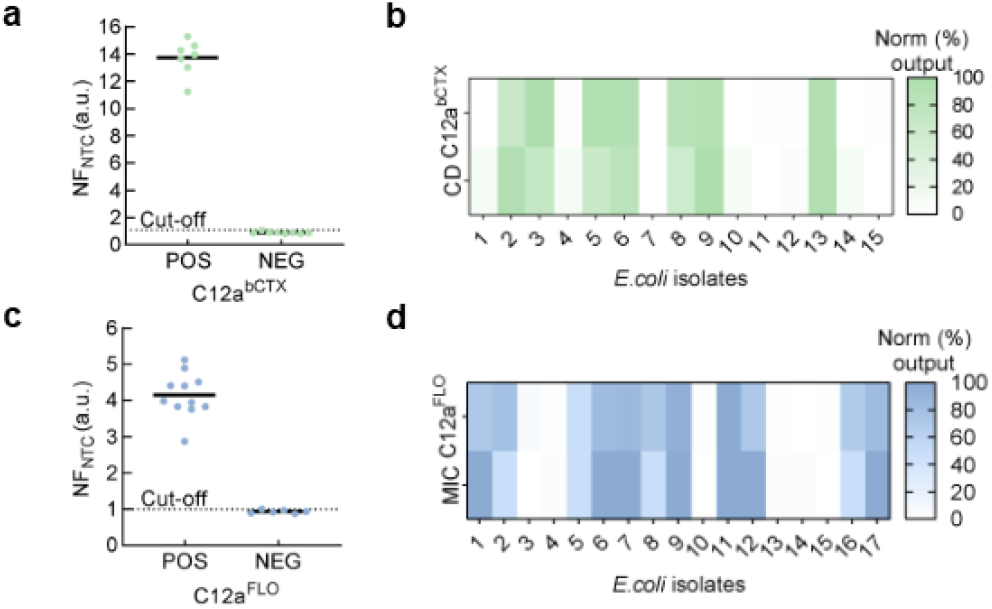
C12a^bCTX^ and C12a^FLO^ comparison with AST methods. **(a)** Distribution of *blaCTX-M-*15 in 17 isolates tested using the C12a^bCTX^ assay. The dotted line represents the LoB at 1.1. **(b)** Heat map comparison of the C12a^bCTX^ assay with the disk diffusion test for AMR (CD). Results were normalized to percentages, using the maximum value obtained for each dataset as 100%. **(c)** Similar to (a), but for the C12a^FLO^ assay. The dotted line indicates the LoB at 1.0. **(d)** Similar to (b) but for the C12a^FLO^ assay with microdilution test (MIC), showing that *floR* positive samples were phenotypically resistant to florfenicol.

Out of 15 isolates, 7 were ESBL producers and found positive by C12a^bCTX^ for the presence of *blaCTX-M-15* gene while 8 were susceptible to beta lactams and found negative by the C12a toolbox (Figure 4A). Similarly, C12a^FLO^ found the *floR* gene in 11 florfenicol resistant isolates, while susceptible isolates (6/17 isolates) turned out negatives by C12a (Figure 4C) (Table S4-5). When assessing the performance of the C12a toolbox, NF_NTC_ values for ARG-positive samples ranged from 11 to 16 for C12a^bCTX^ and from 3 to 4.5 for C12a^FLO^ after 20 minutes of incubation, indicating the detection of the target genes. In contrast, ARG-negative samples showed no NF_NTC_ values higher than 1.0 with both C12a assays, indicating the absence of the target genes (Figure 4A-C). Altogether, the noise to signal ratio, for positive samples, shown by the C12a toolbox was between 3-to 11-fold over the LoB. Full concordance was observed between the presence of resistance genes using the C12a toolbox and AST assays (Figure 4B-D).

### Detection of the *intI1* gene by C12a^**INT**^

Using the C12a toolbox, we evaluated the detection of the *intI1* integrase gene, located in the 5′ conserved region of the Class 1 integron. Previous studies suggested a correlation between *intI1* presence and MDR profiles [32], [33]. Here, we evaluated the detection of the *intI1* gene by designing a set of crRNAs (C12a^INT^) and using a reported set of primers [42] (Table S2) to detect a 146 bp amplification product (Figure S7). Among the three candidate crRNAs, crRNA^int1c^ showed better NF_NTC_ values at 20 minutes reaction time under standardized C12a reaction conditions (Figure 5A). Using the C12a^INT^, we detected the presence of the *intI1* gene in 17 out of 18 ARG-positive samples (Figure 5C). Fluorescence ratios over the NTC control ranged between 3 to 10, showing greater dispersion if compared to the other C12a tools (Figure 5B). Thus, the results could suggest that the presence of *intI1* is associated to *blaCTX-M-15* or *floR*. However, several reports indicate that none of the evaluated ARG are usually located within the gene cassette for integron 1. To get further insights, we investigated the correlation by *in silico* analysis of sequencing data and characterize the integron structure and ARG content within cassette arrays (Figure 5C). Data analysis of short reads using *blaCTX-M15* or *floR* positive *E. coli* isolates revealed that 11/18 carried a cassette array (61.1 %), indicated by the presence of the attC recombination sites. The cassette array frequently contained genes conferring resistance to aminoglycosides (100%), trimethoprim (81.82%), phenicols (36.36%), lincosamide (27.27%) and sulfonamides (18.18%). Additionally, genes conferring tolerance to disinfectants such as *qacE* were observed in 5 isolates (45.45%) (Figure 5C). The *in-silico* analysis revealed that transposons localized near Class 1 Integron were probably involved in the mechanism of acquisition and transfer of the observed cassette array. Consistent with previous reports, *blaCTX-M-15* or *floR* genes were not found as part of the cassette arrays (Table S7). Our results suggest the potential of C12a^INT^ for Class 1 integron detection as proxy of multidrug resistance as previously described [32], [33], however, direct ARG detection remains necessary for surveillance of specific genetic markers of antibiotic resistance.

**Figure 5.**
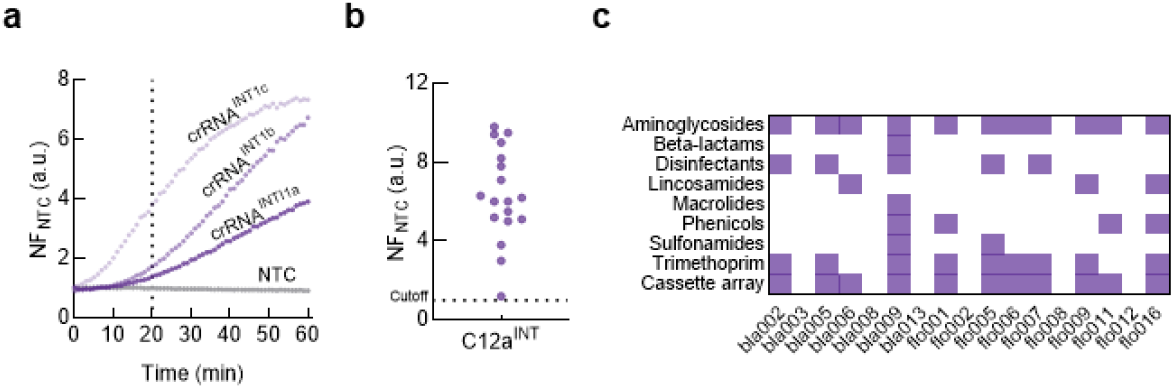
C12a^INT^ detects the *intI1* gene, as potential indicator of multidrug resistant *E. coli*. **(A)** Time-course of normalized fluorescence development for the C12a^INT^ assay with three crRNA designed to detect the *intI1* gene. The vertical dotted line represents the incubation time used. **(B)** CRISPR-Cas detection of *intI1* gene in 18 ARG-positive isolates tested using the C12a^INT^ assay. The dotted line represents the LoB at 1.0. **(C)** Heat map summarizing the antibiotic resistance genes related to different antibiotic families identified within the Class 1 integron cassette array of the ARG-positive *E. coli* isolates used in this study. ARGs were identified by bioinformatic analysis of genome sequencing data (BioProject number PRJNA821865).

### Efficient C12a-mediated ARGs detection in different laboratory setups

Differences in equipment availability may impact on the analytical performance of the C12a detection toolbox. To address this, we adapted our detection assays to three laboratory setups: i) Low-equipped with minimal tools for qualitative readouts, such as a thermocycler and a transilluminator; ii) portable setup using the Lateral Flow Assay (LFA) and transportable workstation (BentoLab^Ⓡ^) suitable for on-site analysis, and iii) High equipped with a multimode microplate reader for quantitative analysis, as used here for assay validation (Figure 6).

**Figure 6.**
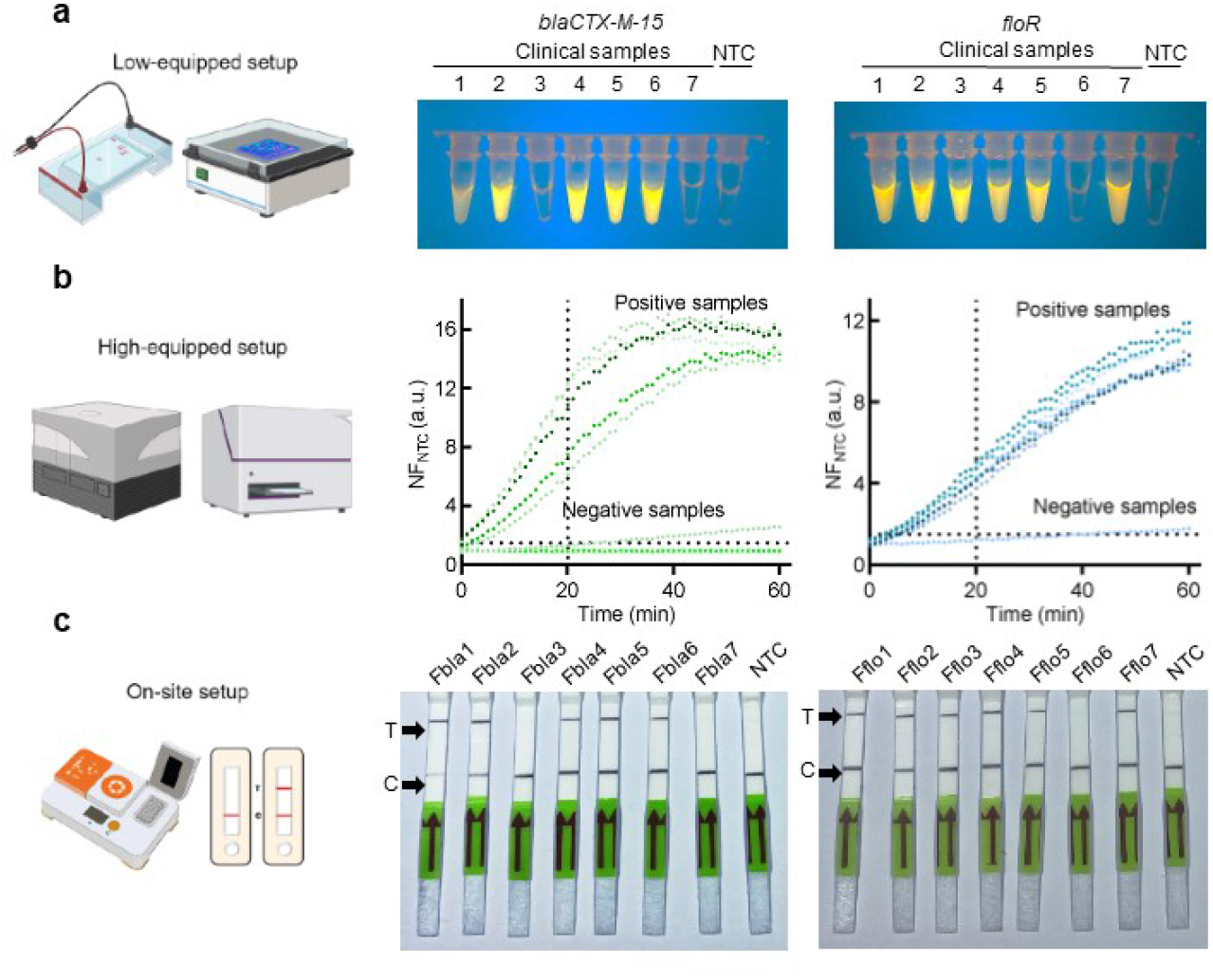
C12a application across different laboratory setups. **(A)** *blaCTX-M-15* and *floR* detection in a low-equipped laboratory setup. Fluorescence was observed by direct visualization and photographed with a smartphone camera, default settings using a blue light transilluminator. **(B)** Similar to (A), but for a high-equipped laboratory. NF_NTC_ (a.u.) values were measured over 60 minutes with a 20-minute detection cutoff. The horizontal dotted line represents an average NF_NTC_ cut-off value of 1.0. **(C)** Similar to (A), but for an on-site setup. Detection by LFA strips readouts at 15 minutes incubation, where the T bands indicate the positivity of the test and C bands correspond to the control. NTC refers to non-template control.

Fourteen purified total fecal DNA available from the stool sample repository (BioProject number PRJNA821865) were analyzed with the C12a^bCTX^ and C12a^FLO^ assays across the three laboratory set-ups. For low equipped setup, direct visualization using a blue light transilluminator showed fluorescence in five out of 7 samples for *blaCTX-M-15*. For *floR*, six out of seven samples resulted in fluorescence observation (Figure 6A). The high-equipped setup showed that C12a^bCTX^ detected *blaCTX-M-15* in five samples, and two samples tested negative, respectively. All positive samples showed NF_NTC_ (a.u.) values exceeding 6 within 20 minutes. Similarly, for the C12a^FLO^, NF_NTC_ (a.u.) values surpassing 3 were recorded at 20 minutes (Figure 6B). For on-site setups, detection relied on LFA readouts instead of fluorescence detection. Here, a double-labeled probe with FITC and biotin is added in the CRISPR-Cas system allowing the appearance of two bands (the control line and test line) on the strip when an ARG is detected (Figure 6C). No difference in detection performance was observed across all setups evaluated. Although the sample size was limited, these results suggest that the C12a toolbox demonstrates sufficient analytical sensitivity and robustness for ARG detection in total DNA samples across different laboratory setups.

## Discussion

The global challenge of antimicrobial resistance (AMR) needs the development of new molecular detection strategies that are rapid, sensitive, cost-effective, and adaptable to diverse contexts, including clinical, veterinary, and environmental settings. While established platforms such as antimicrobial susceptibility testing (AST), PCR, and whole-genome sequencing (WGS) are solid approaches, they remain limited by infrastructure requirements, cost, or turnaround time [34], [35]. In this study, we presented the proof-of-concept of C12a for detecting key antibiotic resistance genes (ARGs) and associated elements. Although not yet validated for clinical or diagnostic use, the C12a toolbox demonstrates promising analytical features that could support future diagnostic development and surveillance efforts.

Molecular detection by C12a display distinctive features: rapid (20 min detection, 100 minutes total analysis), competitive low limits of detection, and support for multiple readout formats, that make it suitable for various applications. CRISPR-Cas-based assays have demonstrated analytical performance comparable to well established techniques, including qPCR, which remains the reference method in most molecular detection workflows [43]. Previous studies showed complete concordance between CRISPR-Cas and qPCR results for the detection of genetic elements in pathogenic and virulence targets [44], [45], [46], [47], [48]. The high specificity of C12a systems aligns with other CRISPR-Cas assays thanks to the dual recognition steps mechanism, which involves both primer selection and crRNA binding. While a direct comparison with qPCR was not conducted in this study, the C12a system showed strong signal-to-noise ratios, well-defined limits of blank (LoB), low limits of detection (LoD), and full concordance with both AST and WGS analysis, reinforcing the reliability of the detection setup. Additionally, the LoD obtained for the C12a toolbox is consistent with those reported in other CRISPR-based studies [43], [49].

The targets selected for this study address both diagnostic relevance and ecological concern. Detection of *blaCTX-M-15* and *floR* genes are of increasing importance due to their widespread distribution across human, animal and environmental interfaces[10] (Figure 1A). Specifically, *blaCTX-M-15*[20], [21] has been traditionally associated with humans and *floR*[15], [18], [19] with animals, however recent findings indicate that both genes are now being detected across diverse scenarios. Resistance gene *floR*, mainly linked to animal populations, has been recently identified in humans, highlighting its spread beyond conventional boundaries [15]. These patterns of occurrence reflect the interconnected dynamics of AMR, where elements like Class 1 integrons, often inserted in transposons and plasmids, contribute to the spread of multidrug-resistant genes in various environments[33]. Although our analysis revealed no integron association for *blaCTX-M-15* or *floR* in the tested isolates, the frequent co-occurrence of *intI1* with other ARGs highlights its utility as a marker of anthropogenic AMR pressure[25], [28], [32], [33].

A key feature of the C12a toolbox is its adaptability to laboratories with varying levels of infrastructure. C12a was successfully used with high-end equipment settings using fluorescence microplate readers, in low-tech settings using basic fluorescence visualization, and in portable formats using lateral flow assays (LFA). Performance was consistent across platforms, with only modest reductions in sensitivity observed in low-resource setups. These results suggest that the C12a system could support surveillance workflows in resource-limited environments, although further testing and optimization for varying sample types (e.g., water, soil) and including other bacterial species are needed. Furthermore, while the system offers a strong biochemical and analytical foundation, clinical diagnostic application will require rigorous validation as by regulatory guidelines. Integration with mobile data platforms may also enhance usability for surveillance or epidemiological purposes.

In summary, this work presents a validated proof-of-concept for a CRISPR-Cas12a-based molecular detection system targeting AMR gene markers. Although not yet suitable for diagnostic implementation, the C12a toolbox demonstrates key features (velocity, sensitivity, specificity, and versatility) that are essential for future development of low-cost, field-adaptable AMR surveillance tools within a One Health framework.

## Materials and Methods

### Biological samples

Two types of biological samples were used to evaluate the performance and adaptability of the C12a toolbox: (i) Total DNA extracted from *Escherichia coli* isolates obtained from fecal samples and (ii) total DNA extracted directly from fecal material. All samples originated from a previously conducted community-based study involving children in Lima, Peru (NIH R01AI108695-01A1) [50], which received ethical approval from the institutional review board at Universidad Peruana Cayetano Heredia (UPCH). No demographic or identifying metadata from participants were accessed for this study. A subset of positive and negative isolates for *blaCTX-M-15* (bla001 – bla015, Table S4) and *flor* (flo001 – flo0017, Table S5) was randomly selected for the study. Only *E. coli* isolates positive for *blaCTX-M-15* or *floR* were tested for the Class 1 integron detection. Total fecal DNA from a subset of samples (Fbla1 – Fbla7, and Fflo1 – Fflo7; Table S6) was used to evaluate the adaptability of the CRISPR assays across three laboratory configurations, including high-, low-, and portable-equipment settings.

### Molecular stocks

#### DNA purification

Total DNA was extracted from an *E. coli* culture grown to OD_600_ = 0.7 using the Quick DNA Miniprep kit (Cat # D4069, Zymo Research) following the manufacturer’s protocol. DNA was quantified using a NanoDrop spectrophotometer and stored at −20°C. Amplicons were purified using the Oligo Clean Kit (Cat # D4060, Zymo Research). *Oligonucleotides synthesis:* Primers and crRNA DNA sequences were sourced from Macrogen Inc. (Seoul, South Korea). *crRNA synthesis:* crRNA was synthesized by *in vitro* transcription using the TranscriptAid T7 High Yield Transcription Kit (Cat#K0411, ThermoFisher Scientific) and purified with the RNA Clean & Concentrator Kit (Cat #11-353B, Zymo Research). *Enzymes:* Taq DNA polymerase and Cas12a proteins were sourced from locally expressed and purified stocks stored at −80°C, as detailed in [51].

### Antimicrobial susceptibility test

#### Combined disk test

The antibiotic susceptibility profile of 32 *E. coli* isolates was determined by the disk diffusion method, using *E. coli* ATCC 25922 (Cat # 0335, Lab Elite) as the control strain. *E. coli* isolates from glycerol stocks were streaked on LB agar plates and incubated overnight at 37°C. Isolated colonies were resuspended in 0.9% saline solution to a density of 0.5 McFarland and swabbed onto Mueller-Hinton agar plates (Cat # M173-500G, Himedia). Antibiotic disks, including cefotaxime (CTX) 30 µg (Cat # 9017, Liofilchem), and ceftazidime (CAZ) 30 µg (Cat # 9019, Liofilchem) were placed on the surface of the medium. After 18 hours of incubation at 37°, inhibition zones were measured and interpreted following the CLSI guidelines. A confirmatory test for ESBL production, recommended by CLSI[52], was performed using CTX 30 µg and CAZ 30 µg disks combined with clavulanic acid: CTX/AC 40 µg (Cat # 9182, Liofilchem), and CAZ/AC 40 µg (Cat # 9145, Liofilchem). Plates were incubated for 18 hours at 37°C, and ESBL producers were detected by measuring the difference in the inhibition halo of the antibiotic in the presence of clavulanic acid (>5 mm). *Broth microdilution assay*: Florfenicol (Cat #F1427-500MG, Merck) was dissolved in dimethyl sulfoxide (DMSO) to a concentration of 100 mg/ml. Serial dilutions, ranging from 256 µg/ml to 0.125 µg/ml were prepared in a 96-well plate (Cat # 3596, ThermoFisher) using Mueller-Hinton Broth II (Cat # 610218, Liofilchem). Each well was inoculated with 1.5 × 10^6^ CFU/ml of cells. After 18 h of incubation at 37°C, the MIC was determined according to CLSI guidelines[53]. A MIC value of 32 µg/mL has been proposed as a tentative threshold to identify florfenicol-resistant *E. coli* isolates harboring the *floR* gene [54]. However, it does not represent a formally established clinical breakpoint.

### Bioinformatic analysis

To identify detection targets, all available *E. coli* resistance gene sequences for *blaCTX-M-15* and *floR* in GenBank as of January 2022 were collected. Multiple alignment was performed, obtaining a consensus sequence of 1057 bp for *blaCTX-M-15* and 1415 bp for *floR* (Supplementary Dataset 1 and Dataset 2, respectively). Primer sets were designed for the *blaCTX-M-15* [16] and *floR* [19]genes. For crRNA guide design and evaluation, CRISPRscan and Chop-Chop bioinformatics tools were used [55], [56]. DNA data analysis was downloaded from the BioProject web portal of the NCBI (https://www.ncbi.nlm.nih.gov/bioproject/) using accession code PRJNA821865. ARG presence was evaluated using the ResFinder platform (http://genepi.food.dtu.dk/resfinder) v4.6.0 [57] with default parameters. Briefly, a minimum length of 60% and a threshold of 90% were set up for ARG and disinfectant-resistance gene identification. *E*.*coli* was defined as the species input. For the identification of the integrase-integron class 1, the VRprofile2 platform (https://tool2-mml.sjtu.edu.cn/VRprofile/)[58] with default parameters. Fluorescence data from the Synergy H1 plate reader were collected using Gen 5 software and exported to Excel. NF_ntc_ values were calculated during data processing and analysis as appropriate. All fluorescence data analysis and chart preparation were performed using GraphPad Prism version 10.0.0, www.graphpad.com.

### C12a Molecular Assays

#### DNA amplification and analysis

PCR conditions for amplifying the *blaCTX-M-15* and *floR* targets included an initial denaturation at 95°C for 3 minutes, followed by 30 cycles of denaturation at 95° for 30 sec, annealing at 59°C for 30 sec, extension at 72°C for 30 sec, and a final extension at 72°C for 5 minutes. For the *intI1* gene target, an annealing temperature of 60°C was used. Amplification reactions (20 µL) contained 2 ng/µL Taq DNA polymerase, 0.2 µM primers, and 1 to 10 ng/µl of DNA in RPB1X buffer (Table S1). PCR products were visualized using 1.7% agarose gel electrophoresis (Cat # CSL-AG500, Cleaver Scientific) in 1X TBE buffer (Table S1) at 80 V for 1 hour. Gels were stained with 6X Trick-Track – SYBR Gold (Cat #R1161, ThermoFisher) and visualized with a Gel Doc system (Bio-Rad) using the Sybergold option. A 100 bp molecular weight marker (Cat #SM0242, Thermo Scientific) was included. Band intensity was quantified as AUC using Gel Analyzer v23 (www.gelanalyzer.com).

#### crRNA/Cas12a-mediated detection

150 nM crRNA was refolded (65°C for 10 minutes, followed by 10 minutes at 25°C) and assembled with 100 nM Cas12a, and 2 µM FAM probe (6-FAM/TTATT/3IABkFQ) in CRB1 buffer (Table S1) in the dark for 10 minutes. Unpurified PCR reactions containing amplicons (5 µL) were combined with 10 µL of the complex in CRB2 buffer (Table S1) to a final volume of 100 µL. Fluorescence was measured using a plate reader (Synergy H1, BioTek Instruments) with excitation at 491 nm and emission at 525 nm.

#### Lateral Flow Assay (LFA)

CRISPR-Cas reaction was performed as described above using a biotin/fluorescein probe (6-FAM/TTATTATT/BIOTIN). The reaction buffer was supplemented with polyethylene glycol (Cat # 25322-68-3, Merck) to a final concentration of 5%. After 30 minutes of incubation, a LFA strip (Cat # 3822-9000, Milenia Biotec GmbH) was submerged in the reaction. The signal on the strips was detected after 5-10 minutes.

#### LoB and LoD determination

Pure genomic DNA from ten susceptible *E. coli* isolates (true negative) were used to determine the LoB, using the formula LoB = Avg_NF_NTC_ + 2 SD (where “Avg” is average NF_ntc_ values, and “SD” is standard deviation). The LoD was determined using both amplification products and bacterial cell counts for the *blaCTX-M-15* and *floR* targets. Amplicons were purified using the Oligo Clean Kit (Cat # D4060, Zymo Research) and quantified by absorbance at 260nm. DNA concentrations ranged from 1 aM to 10 nM. The LoD was then estimated as the lowest DNA concentration above the LoB + 2SD [59]. For the bacterial cell LoD determination, a curve was generated using a positive *E. coli* isolate culture. Starting from a cell density of 0.5 McFarland, and 7 ten-fold dilutions in duplicate. One set of dilutions was plated on LB agar plates for CFU counting, while the other set was analyzed directly with C12a.

## Supporting information

Sequences

## Ethics

The *E. coli* isolates and DNA fecal samples used in this study were anonymous and derived from a biorepository of a longitudinal study approved by the ethics committee of Universidad Peruana de Cayetano Heredia (UPCH, SIDISI: 65178).

## Authors Contribution

M.V-R., R.A., M.P., and P.M. conceived the project. M.V-R., R.A., and P.M. designed experiments. M.V-R., R.A., and S.A. performed experiments. M.V-R., R.A., and P.M. analyzed the data. M.V-R., R.A., K.P., and P.M. elaborated figures and tables. R.S. and D.P. helped with data interpretation. M.V-R., R.A., S.A., K.P., and P.M. wrote the manuscript with the input of D.P., R.S., and M.P.

## Acknowledgments

We are very thankful to Dr. Maya Nadimpalli for promptly providing sequencing data of the *E. coli* isolate collection used in this study.

## References

[1] Hutchings, M. I., Truman, A. W. & Wilkinson, B. Antibiotics: past, present and future. Curr Opin Microbiol 51, 72–80 (2019).

[2] Lipsitch, M. & Samore, M. H. Antimicrobial Use and Antimicrobial Resistance: A Population Perspective. Emerg Infect Dis 8, 347–354 (2002).

[3] Bungau, S., Tit, D. M., Behl, T., Aleya, L. & Zaha, D. C. Aspects of excessive antibiotic consumption and environmental influences correlated with the occurrence of resistance to antimicrobial agents. Curr Opin Environ Sci Health 19, 100224 (2021).

[4] Huijbers, P. M. C. et al. Role of the Environment in the Transmission of Antimicrobial Resistance to Humans: A Review. Environ Sci Technol 49, 11993–12004 (2015).

[5] Zhang, Z. et al. Assessment of global health risk of antibiotic resistance genes. Nat Commun 13, 1553 (2022).

[6] von Wintersdorff, C. J. H. et al. Dissemination of Antimicrobial Resistance in Microbial Ecosystems through Horizontal Gene Transfer. Front Microbiol 7, (2016).

[7] Arnold, K. E. et al. The need for One Health systems-thinking approaches to understand multiscale dissemination of antimicrobial resistance. Lancet Planet Health 8, e124–e133 (2024).

[8] Samreen Ahmad, I., Malak, H. A. & Abulreesh, H. H. Environmental antimicrobial resistance and its drivers: a potential threat to public health. J Glob Antimicrob Resist 27, 101–111 (2021).

[9] Aslam, B. et al. Antibiotic Resistance: One Health One World Outlook. Front Cell Infect Microbiol 11, (2021).

[10] Zhuang, M. et al. Distribution of antibiotic resistance genes in the environment. Environmental Pollution 285, 117402 (2021).

[11] Mazumder, R., Abdullah, A., Ahmed, D. & Hussain, A. High Prevalence of blaCTX-M-15 Gene among Extended-Spectrum β-Lactamase-Producing Escherichia coli Isolates Causing Extraintestinal Infections in Bangladesh. Antibiotics 9, 796 (2020).

[12] Akenten, C. W. et al. Carriage of ESBL-producing Klebsiella pneumoniae and Escherichia coli among children in rural Ghana: a cross-sectional study. Antimicrob Resist Infect Control 12, 60 (2023).

[13] Li, P. et al. Analysis of Resistance to Florfenicol and the Related Mechanism of Dissemination in Different Animal-Derived Bacteria. Front Cell Infect Microbiol 10, (2020).

[14] Ying, Y. et al. Florfenicol Resistance in Enterobacteriaceae and Whole-Genome Sequence Analysis of Florfenicol-Resistant Leclercia adecarboxylata Strain R25. Int J Genomics 2019, 1–10 (2019).

[15] Murray, M. et al. Market Chickens as a Source of Antibiotic-Resistant Escherichia coli in a Peri-Urban Community in Lima, Peru. Front Microbiol 12, (2021).

[16] Ramadan, A. A., Abdelaziz, N. A., Amin, M. A. & Aziz, R. K. Novel blaCTX-M variants and genotype-phenotype correlations among clinical isolates of extended spectrum beta lactamase-producing Escherichia coli. Sci Rep 9, 4224 (2019).

[17] Negeri, A. A. et al. Characterization of plasmids carrying blaCTX-M genes among extra-intestinal Escherichia coli clinical isolates in Ethiopia. Sci Rep 13, 8595 (2023).

[18] Lu, J. et al. Spread of the florfenicol resistance floR gene among clinical Klebsiella pneumoniae isolates in China. Antimicrob Resist Infect Control 7, 127 (2018).

[19] Qian, C. et al. Identification of floR Variants Associated With a Novel Tn4371-Like Integrative and Conjugative Element in Clinical Pseudomonas aeruginosa Isolates. Front Cell Infect Microbiol 11, (2021).

[20] Gundran, R. S. et al. Prevalence and distribution of blaCTX-M, blaSHV, blaTEM genes in extended-spectrum β-lactamase-producing E. coli isolates from broiler farms in the Philippines. BMC Vet Res 15, 227 (2019).

[21] Higgins, O. et al. Portable Differential Detection of CTX-M ESBL Gene Variants, bla _CTX-M-1_ and bla _CTX-M-15_, from Escherichia coli Isolates and Animal Fecal Samples Using Loop-Primer Endonuclease Cleavage Loop-Mediated Isothermal Amplification. Microbiol Spectr 11, (2023).

[22] Rangama, S. et al. Mechanisms Involved in the Active Secretion of CTX-M-15 β-Lactamase by Pathogenic Escherichia coli ST131. Antimicrob Agents Chemother 65, (2021).

[23] Liao, C.-Y. et al. Antimicrobial Resistance of Escherichia coli From Aquaculture Farms and Their Environment in Zhanjiang, China. Front Vet Sci 8, (2021).

[24] Tokuda, M. & Shintani, M. Microbial evolution through horizontal gene transfer by mobile genetic elements. Microb Biotechnol 17, (2024).

[25] Gillings, M. R. et al. Using the class 1 integron-integrase gene as a proxy for anthropogenic pollution. ISME J 9, 1269–1279 (2015).

[26] Zhang, S. et al. Dissemination of antibiotic resistance genes (ARGs) via integrons in Escherichia coli: A risk to human health. Environmental Pollution 266, 115260 (2020).

[27] Bhat, B. A. et al. Integrons in the development of antimicrobial resistance: critical review and perspectives. Front Microbiol 14, (2023).

[28] Corno, G. et al. Class 1 integron and related antimicrobial resistance gene dynamics along a complex freshwater system affected by different anthropogenic pressures. Environmental Pollution 316, 120601 (2023).

[29] Stalder, T., Barraud, O., Casellas, M., Dagot, C. & Ploy, M.-C. Integron Involvement in Environmental Spread of Antibiotic Resistance. Front Microbiol 3, (2012).

[30] Halaji, M. et al. The Global Prevalence of Class 1 Integron and Associated Antibiotic Resistance in Escherichia coli from Patients with Urinary Tract Infections, a Systematic Review and Meta-Analysis. Microbial Drug Resistance 26, 1208–1218 (2020).

[31] Stedtfeld, R. D. et al. Isothermal assay targeting class 1 integrase gene for environmental surveillance of antibiotic resistance markers. J Environ Manage 198, 213–220 (2017).

[32] Rao, S., Maddox, C. W., Hoien-Dalen, P., Lanka, S. & Weigel, R. M. Diagnostic Accuracy of Class 1 Integron PCR Method in Detection of Antibiotic Resistance in Salmonella Isolates from Swine Production Systems. J Clin Microbiol 46, 916–920 (2008).

[33] Solis, M. N. et al. Detecting Class 1 Integrons and Their Variable Regions in Escherichia coli Whole-Genome Sequences Reported from Andean Community Countries. Antibiotics 13, 394 (2024).

[34] Abramova, A., Berendonk, T. U. & Bengtsson-Palme, J. A global baseline for qPCR-determined antimicrobial resistance gene prevalence across environments. Environ Int 178, 108084 (2023).

[35] Abramova, A., Karkman, A. & Bengtsson-Palme, J. Metagenomic assemblies tend to break around antibiotic resistance genes. BMC Genomics 25, 959 (2024).

[36] Li, S.-Y. et al. CRISPR-Cas12a-assisted nucleic acid detection. Cell Discov 4, 20 (2018).

[37] Chen, J. S. et al. CRISPR-Cas12a target binding unleashes indiscriminate single-stranded DNase activity. Science (1979) 360, 436–439 (2018).

[38] Broughton, J. P. et al. CRISPR–Cas12-based detection of SARS-CoV-2. Nat Biotechnol 38, 870–874 (2020).

[39] Karimi Dehkordi, M., Halaji, M. & Nouri, S. Prevalence of class 1 integron in Escherichia coli isolated from animal sources in Iran: a systematic review and meta-analysis. Trop Med Health 48, 16 (2020).

[40] Fadare, F. T., Fadare, T. O. & Okoh, A. I. Prevalence, molecular characterization of integrons and its associated gene cassettes in Klebsiella pneumoniae and K. oxytoca recovered from diverse environmental matrices. Sci Rep 13, 14373 (2023).

[41] Nadimpalli, M. L. et al. Effects of breastfeeding on children’s gut colonization with multidrug-resistant Enterobacterales in peri-urban Lima, Peru. Gut Microbes 16, (2024).

[42] Zhang, Y., Lu, J., Wu, J., Wang, J. & Luo, Y. Potential risks of microplastics combined with superbugs: Enrichment of antibiotic resistant bacteria on the surface of microplastics in mariculture system. Ecotoxicol Environ Saf 187, 109852 (2020).

[43] Kaminski, M. M., Abudayyeh, O. O., Gootenberg, J. S., Zhang, F. & Collins, J. J. CRISPR-based diagnostics. Nat Biomed Eng 5, 643–656 (2021).

[44] Wang, B. et al. Cas12aVDet: A CRISPR/Cas12a-Based Platform for Rapid and Visual Nucleic Acid Detection. Anal Chem 91, 12156–12161 (2019).

[45] Curti, L. A. et al. CRISPR-based platform for carbapenemases and emerging viruses detection using Cas12a (Cpf1) effector nuclease. Emerg Microbes Infect 9, 1140–1148 (2020).

[46] Wang, X. et al. Development and clinical application of a novel CRISPR-Cas12a based assay for the detection of African swine fever virus. BMC Microbiol 20, 282 (2020).

[47] Wang, D. et al. A CRISPR/Cas12a-based DNAzyme visualization system for rapid, non-electrically dependent detection of Bacillus anthracis. Emerg Microbes Infect 11, 429–438 (2022).

[48] van Dongen, J. E. & Segerink, L. I. Building the Future of Clinical Diagnostics: An Analysis of Potential Benefits and Current Barriers in CRISPR/Cas Diagnostics. ACS Synth Biol 14, 323–331 (2025).

[49] Zhou, J., Li, Z., Seun Olajide, J. & Wang, G. CRISPR/Cas-based nucleic acid detection strategies: Trends and challenges. Heliyon 10, e26179 (2024).

[50] Pajuelo, M. J. et al. Epidemiology of enterotoxigenic Escherichia coli and impact on the growth of children in the first two years of life in Lima, Peru. Front Public Health 12, (2024).

[51] Mendoza-Rojas, G. et al. A low-cost and open-source protocol to produce key enzymes for molecular detection assays. STAR Protoc 2, 100899 (2021).

[52] CLSI. Performance Standards for Antimicrobial Susceptibility Testing; Twenty-First Informational Supplement M100-S21. (Wayne, PA, 2020).

[53] CLSI. Methods for Dilution Antimicrobial Susceptibility Tests for Bacteria That Grow Aerobically M07-Ed12. (Wayne, PA, 2024).

[54] Singer, R. S. et al. Relationship between Phenotypic and Genotypic Florfenicol Resistance in Escherichia coli. Antimicrob Agents Chemother 48, 4047–4049 (2004).

[55] Labun, K. et al. CHOPCHOP v3: expanding the CRISPR web toolbox beyond genome editing. Nucleic Acids Res 47, W171–W174 (2019).

[56] Moreno-Mateos, M. A. et al. CRISPRscan: designing highly efficient sgRNAs for CRISPR-Cas9 targeting in vivo. Nat Methods 12, 982–988 (2015).

[57] Bortolaia, V. et al. ResFinder 4.0 for predictions of phenotypes from genotypes. Journal of Antimicrobial Chemotherapy 75, 3491–3500 (2020).

[58] Wang, M. et al. VRprofile2: detection of antibiotic resistance-associated mobilome in bacterial pathogens. Nucleic Acids Res 50, W768–W773 (2022).

[59] CLSI. Evaluation of Detection Capability for Clinical Laboratory Measurement Procedures; Approved Guideline. CLSI Document EP17-A2. (Clinical and Laboratory Standards Institute, Wayne, PA, 2012).

