## Supplementary Figures for "Versatile and Portable Cas12a-mediated Detection of Antibiotic Resistance Markers"

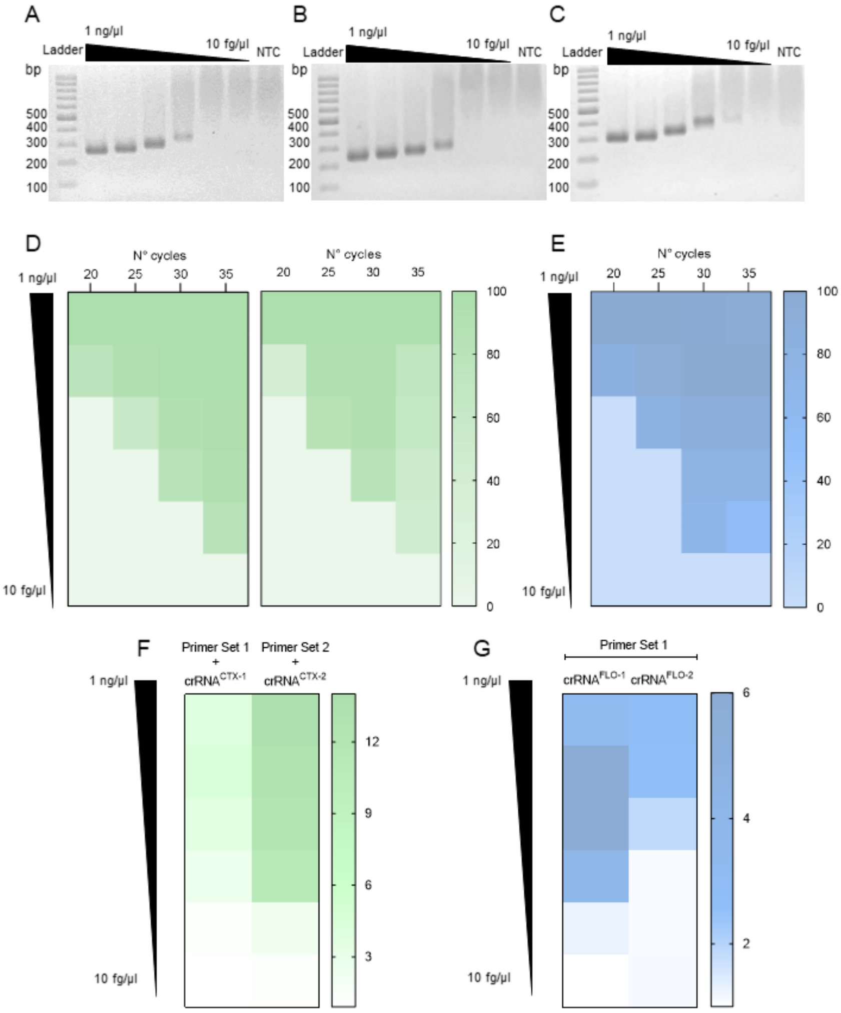


**Supplementary Figure 1. Overview of the optimization process for the CRISPR/Cas12a detection systems. (A), (B),** and **(C)** show agarose gel images illustrating the results of the 30-cycle PCR for the *blaCTX-M-15* primer set 1, primer set 2, and the *floR* primer set, respectively. The observed decrease in migration with reduced fragment concentration is attributed to an interaction with the SYBR Gold dye, as documented in the literature [15]. **(D)** Heatmap displaying the AUC (Area Under the Curve) values of the amplification bands observed in an agarose gel for the *blaCTX-M-15* primer sets. A template titration was conducted in a standard PCR with an increasing number of amplification cycles. **(E)** Similar to (D), but for the *floR* primer set. **(F)** Heatmap displaying fluorescent ratio values for the crRNA candidates designed for the *blaCTX-M-15* gene, obtained from CRISPR-Cas-based detection of 30-cycle PCR amplification products across a template concentration range from 10 fg/µl to 1 ng/µl. **(G)** Similar to (F), but for *floR*.

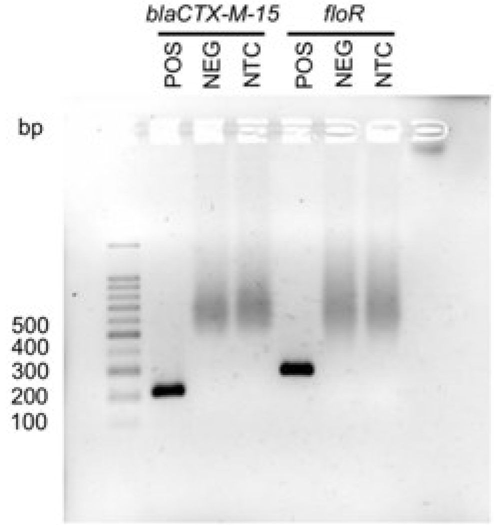


**Supplementary Figure 2. Agarose gel displaying PCR amplification products for the ARG evaluated.** Amplified products obtained from a standard 30-cycle PCR were analyzed using 1.7% agarose gel electrophoresis, with a 100 bp ladder for size reference. Amplicons were observed exclusively in positive samples. POS indicates a reaction containing the target gene; NEG indicates a reaction with template DNA missing the target gene; NTC indicates a reaction without template DNA. Specifically, the primer set 2 targeting the *blaCTX-M-15* gene produced a 216 bp amplicon, while the gene-specific primer for *floR* produced a 270 bp amplicon.

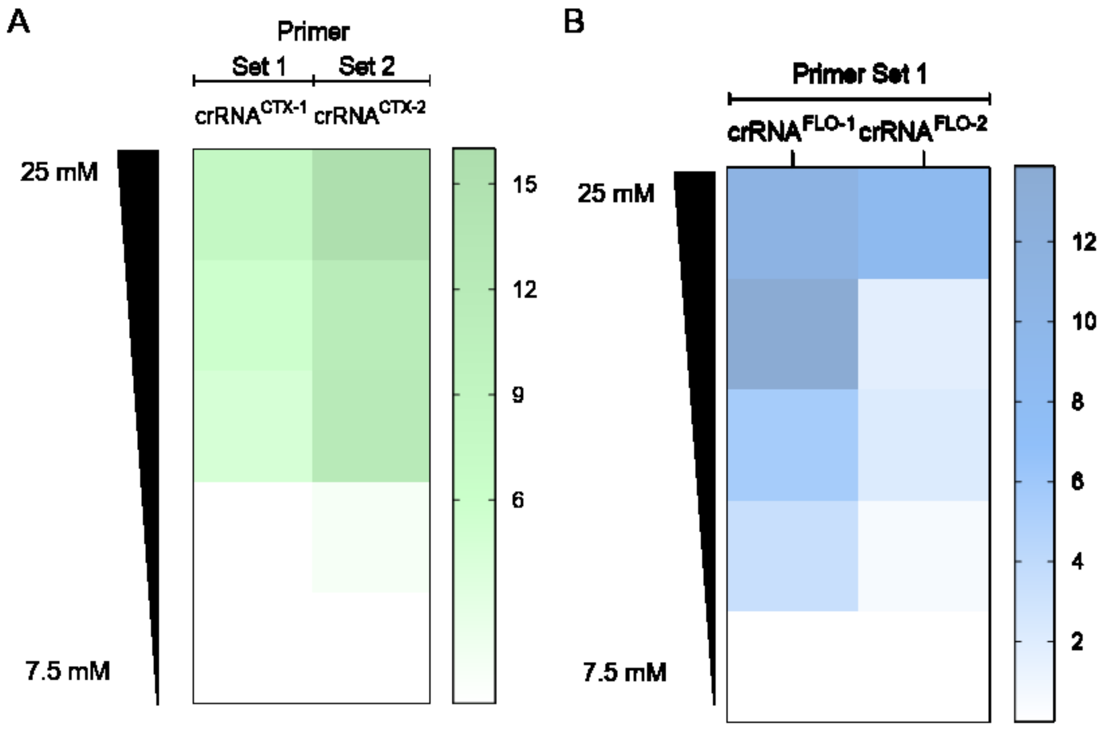


**Supplementary Figure 3. Evaluation of Mg2+ concentration on the CRISPR-Cas system performance.** Heatmaps displaying the $NF_{NTC}$ values of the $MgCl^{2+}$ titration for the crRNA candidates designed for *blaCTX-M-15* and *floR*. The concentration of DNA was 2 ng/µl, meanwhile Mg2+ ranged 7.5 mM to 25 mM. PCR amplicons were obtained using a 30-cycle PCR protocol with primer set 2 for *blaCTX-M-15* and primer set 1 for *floR*. The $NF_{NTC}$ values were measured at 15 min.

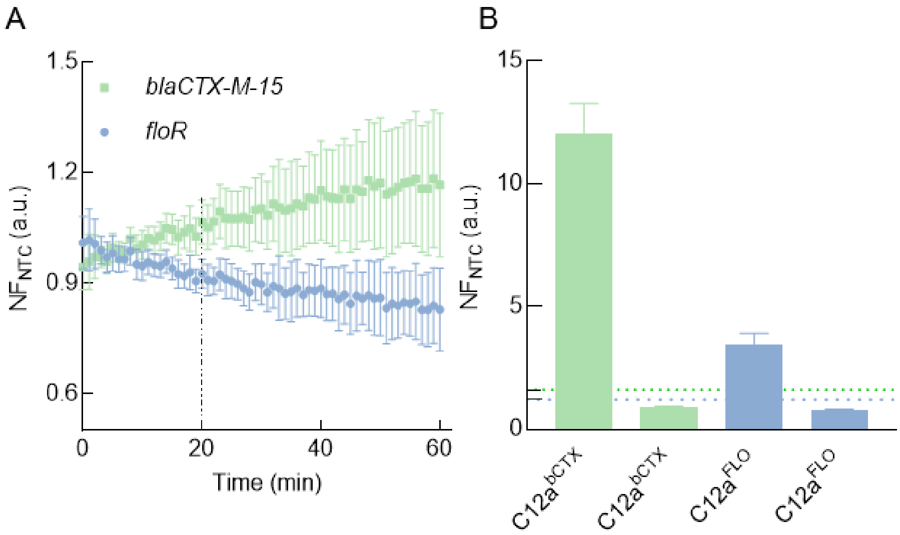


**Supplementary Figure 4. Determination of Limit of Blank (LoB) and Day-to-Day Reproducibility of C12a$^{bCTX}$ and C12a$^{FLO}$ (A)** Normalized fluorescence ratio ($NF_{NTC}$) values over time (minutes) for ten negative samples were analyzed to determine the Limit of Blank (LoB). The average $NF_{NTC}$ values for negative samples after 20 minutes (dotted line) were 1.04 ± 0.02 for *blaCTX-M-15* and 0.94 ± 0.02 for *floR*. Error bars represent the standard deviation of at least ten consecutive measurements. **(B)** $NF_{NTC}$ values for positive and negative samples at 20 minutes detection point across 34 experimental days. The bar represents the mean $NF_{NTC}$ values with the error bars representing the standard deviation. The dotted lines correspond to the LoB values calculated for each gene.

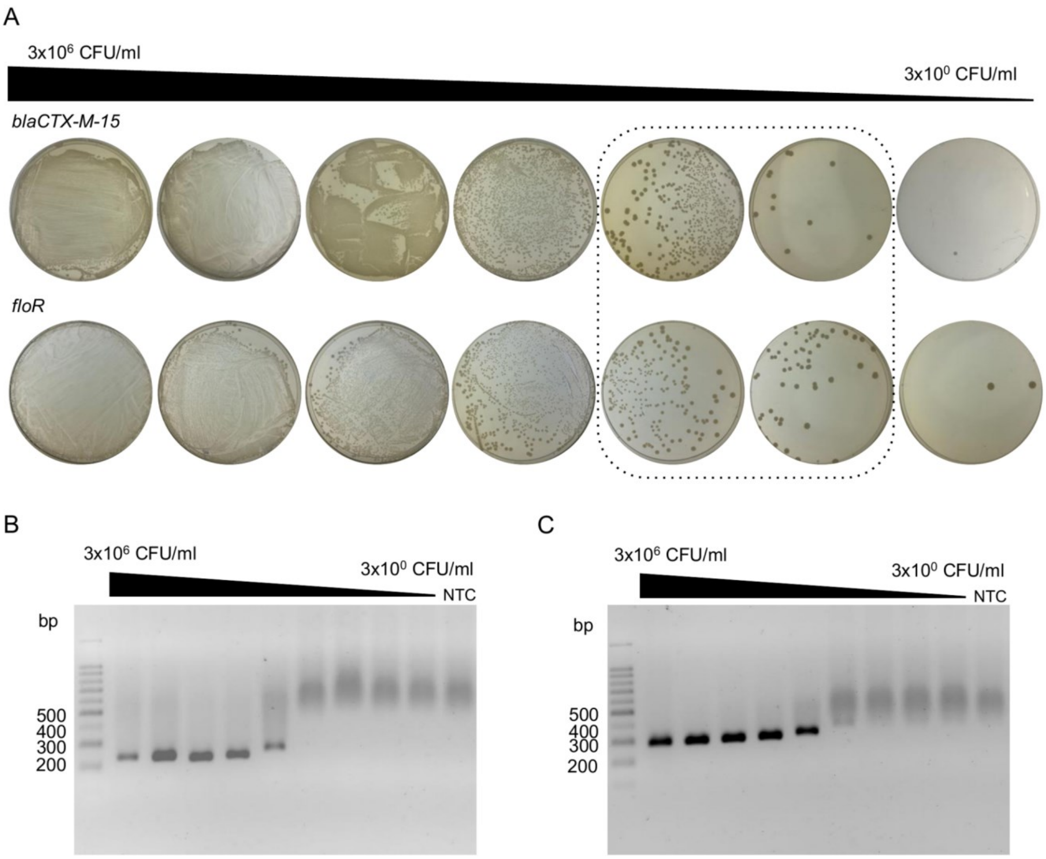


**Supplementary Figure 5. Determination of the Limit of detection based on CFU quantification.** To determine the LoD in CFU/ml, 7 ten-fold dilutions of a cell culture of *E.coli* carrier of the *blaCTX-M-15* and *floR*, respectively at 0.5 Mc Farland were prepared. **(A)** *E.coli* cells were plated on LB agar after 18 hours of incubation at 37°C. The dilution curve was prepared using 1 mL of LB medium, and 50 μL of each dilution was spread on the LB plates. The CFU/ml titer was determined using the CFU number observed in the dilutions framed in the dotted box. **(B)** Amplified products obtained from each dilution using a standard 30-cycle PCR were analyzed using 1.7% agarose gel electrophoresis, with a 100 bp ladder for size comparison. For *blaCTX-M-15* an amplicon of 216 bp was expected. **(C)** Similar to panel (B), but for *floR,* with an expected amplicon size of 270 bp. The reduced migration at lower fragment concentrations is caused by an interaction with the SYBR Gold dye, as documented in the literature [15].

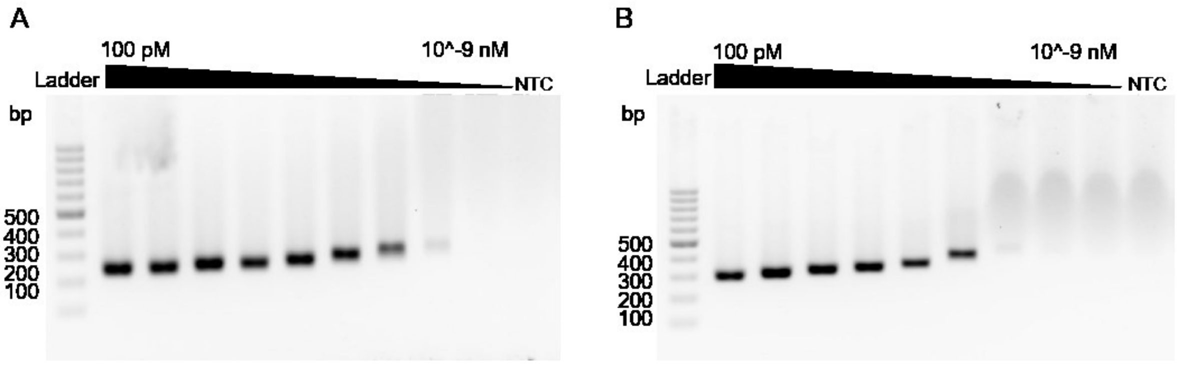


**Supplementary Figure 6. Agarose-gel electrophoresis for comparison of analytical sensitivity between CRISPR-Cas-based fluorescent and naked-eye agarose electrophoresis detection.** Target amplification across different template concentrations over a 30-cycle PCR. Amplified products were resolved using 1.7% agarose gel electrophoresis, alongside a 100 bp ladder for size comparison. **(A)** An amplicon of 216 bp was expected for *blaCTX-M-15.* **(B)** Same as (A), but for *floR* with an expected amplicon size of 270 bp. The observed decrease in migration with reduced fragment concentration is attributed to an interaction with the SYBR Gold dye, as documented in the literature [15].

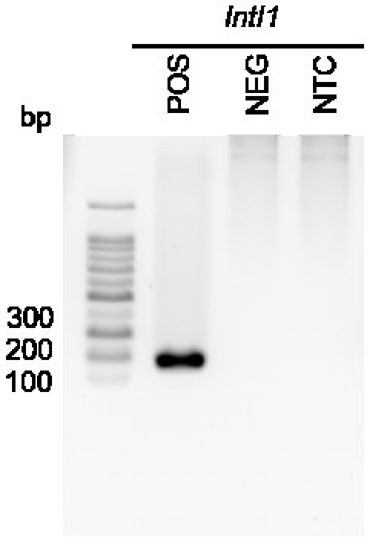


**Supplementary Figure 7. Agarose gel displaying PCR amplification products for the *intI1* gene.** Amplified products obtained from a standard 30-cycle PCR were analyzed using 1.7% agarose gel electrophoresis, with a 100 bp ladder for size reference. Amplicons were observed exclusively in positive samples. Specifically, the primer set targeting the *intI1* gene produced a 146 bp amplicon.

### Supplementary files

**Supplementary Dataset 1. DNA sequences for *blaCTX-M-15* alignment.**

**Supplementary Dataset 2. DNA sequences for *floR* alignment.**

**Supplementary File 1. Excel File with supplementary tables:**

- **Sheet 1:** Supplementary Table 1, List of buffers used in this study.
- **Sheet 2:** Supplementary Table 2, List of oligonucleotides used in this study.
- **Sheet 3:** Supplementary Table 3, List of DNA sequences for *blaCTX-M-15* and *floR* genes amplified with the selected primer sets. DNA sequences were obtained by SANGER sequencing.
- **Sheet 4:** Supplementary Table 4, Database with NGS, CRISPR/Cas, and Antimicrobial Susceptibility Testing data for *E.coli* isolates evaluated for *blaCTX-M-15*.
- **Sheet 5:** Supplementary Table 5, Database with NGS, CRISPR/Cas, and Antimicrobial Susceptibility Testing data for *E.coli* isolates evaluated for *floR*.
- **Sheet 6:** Supplementary Table 6, Database with NGS, CRISPR/Cas, and Antimicrobial Susceptibility Testing data for feces samples.
- **Sheet 7:** Supplementary Table 7, List of ARGs identified in the *blaCTX-M-15* and *floR*-positive *E. coli* isolates, including those found in the genome and in the Class 1 integron.

### References


1. Larsson A. AliView: A fast and lightweight alignment viewer and editor for large datasets. Bioinformatics. 2014 Feb 27;30(22):3276–8.
2. Edgar RC. MUSCLE: Multiple sequence alignment with high accuracy and high throughput. Nucleic Acids Res. 2004;32(5):1792–7.
3. Ramadan AA, Abdelaziz NA, Amin MA, Aziz RK. Novel blaCTX-M variants and genotype-phenotype correlations among clinical isolates of extended spectrum beta lactamase-producing Escherichia coli. Sci Rep. 2019 Dec 1;9(1).
4. Qian C, Liu H, Cao J, Ji Y, Lu W, Lu J, et al. Identification of floR Variants Associated With a Novel Tn4371-Like Integrative and Conjugative Element in Clinical Pseudomonas aeruginosa Isolates. Front Cell Infect Microbiol. 2021 Jun 21;11.
5. Moreno-Mateos MA, Vejnar CE, Beaudoin JD, Fernandez JP, Mis EK, Khokha MK, et al. CRISPRscan: Designing highly efficient sgRNAs for CRISPR-Cas9 targeting in vivo. Nat Methods. 2015 Sep 29;12(10):982–8.
6. Labun K, Montague TG, Krause M, Torres Cleuren YN, Tjeldnes H, Valen E. CHOPCHOP v3: Expanding the CRISPR web toolbox beyond genome editing. Nucleic Acids Res. 2019 Jul 1;47(W1):W171–4.
7. Mathews DH. Predicting RNA secondary structure by free energy minimization. Theor Chem Acc. 2006 Aug;116(1–3):160–8.
8. Mendoza-Rojas, G., Sarabia-Vega, V., Sanchez-Castro, A., Tello, L., Cabrera-Sosa, L., Nakamoto, J. A., Peñaranda, K., Adaui, V., Alcántara, R., & Milón, P. (2021). A low-cost and open-source protocol to produce key enzymes for molecular detection assays. *STAR protocols*, 2(4), 100899. https://doi.org/10.1016/j.xpro.2021.100899
9. CLSI. M100 - Performance standards for antimicrobial susceptibility testing. 2020. 282.
10. CLSI. M07 - Methods for Dilution Antimicrobial Susceptibility Tests for Bacteria That Grow Aerobically - Ed12 [Internet]. 12th ed. 2024. Available from: https://clsi.org/terms-of-use/.
11. Singer RS, Patterson SK, Meier AE, Gibson JK, Lee HL, Maddox CW. Relationship between phenotypic and genotypic florfenicol resistance in Escherichia coli. Antimicrob Agents Chemother.

2004;48(10):4047–9.

12. Bortolaia, V., Kaas, R. S., Ruppe, E., Roberts, M. C., Schwarz, S., Cattoir, V., Philippon, A., Allesoe, R. L., Rebelo, A. R., Florensa, A. F., Fagelhauer, L., Chakraborty, T., Neumann, B., Werner, G., Bender, J. K., Stingl, K., Nguyen, M., Coppens, J., Xavier, B. B., Malhotra-Kumar, S., … Aarestrup, F. M. (2020). ResFinder 4.0 for predictions of phenotypes from genotypes. The Journal of antimicrobial chemotherapy, 75(12), 3491–3500. https://doi.org/10.1093/jac/dkaa345

13. Wang, M., Goh, Y. X., Tai, C., Wang, H., Deng, Z., & Ou, H. Y. (2022). VRprofile2: detection of antibiotic resistance-associated mobilome in bacterial pathogens. Nucleic acids research, 50(W1), W768–W773. https://doi.org/10.1093/nar/gkac321

14. Néron, B., Littner, E., Haudiquet, M., Perrin, A., Cury, J., & Rocha, E. P. C. (2022). IntegronFinder 2.0: Identification and Analysis of Integrons across Bacteria, with a Focus on Antibiotic Resistance in Klebsiella. Microorganisms, 10(4), 700. https://doi.org/10.3390/microorganisms10040700

15. Sharp, P. A., Sugden, B., & Sambrook, J. (1973). Detection of two restriction endonuclease activities in Haemophilus parainfluenzae using analytical agarose--ethidium bromide electrophoresis. Biochemistry, 12(16), 3055–3063. https://doi.org/10.1021/bi00740a018
